# Neural Signatures of Reward and Sensory Prediction Error in Motor Learning

**DOI:** 10.1101/262576

**Authors:** Dimitrios J. Palidis, Joshua G.A. Cashaback, Paul L. Gribble

## Abstract

At least two distinct processes have been identified by which motor commands are adapted according to movement-related feedback: reward based learning and sensory error based learning. In sensory error based learning, mappings between sensory targets and motor commands are recalibrated according to sensory error feedback. In reward based learning, motor commands are associated with subjective value, such that successful actions are reinforced. We designed two tasks to isolate reward and sensory error based motor adaptation, and recorded electroencephalography (EEG) from humans to identify and dissociate the neural correlates of reward and sensory error processing. We designed a visuomotor rotation task to isolate sensory error based learning which was induced by altered visual feedback of hand position. In a reward learning task, we isolated reward based learning induced by binary reward feedback that was decoupled from the visual target. We found that a fronto-central event related potential called the feedback related negativity (FRN) was elicited specifically by reward feedback but not sensory error feedback. A more posterior component called the P300 was evoked by feedback in both tasks. In the visuomotor rotation task, P300 amplitude was increased by sensory error induced by perturbed visual feedback, and was correlated with learning rate. In the reward learning task, P300 amplitude was increased by reward relative to non reward and by surprise regardless of feedback valence. We propose that during motor adaptation, the FRN might specifically mark reward prediction error while the P300 might reflect processing which is modulated more generally by prediction error.

**New and Noteworthy:** We studied the event related potentials evoked by feedback stimuli during motor adaptation tasks that isolate reward and sensory error learning mechanisms. We found that the feedback related negativity was specifically elicited by reward feedback, while the P300 was observed in both tasks. These results reveal neural processes associated with different learning mechanisms and elucidate which classes of errors, from a computational standpoint, elicit the FRN and P300.

## Introduction

It is thought that sensorimotor adaptation is driven by two distinct error signals, sensory prediction error (SPE) and reward prediction error (RPE), and that both can simultaneously contribute to learning (Huang et al. 2011; Izawa and Shadmehr 2011; Shmuelof et al. 2012; Galea et al. 2015; Nikooyan and Ahmed 2015). Electroencephalography (EEG) has been used to identify neural signatures of error processing in various motor learning and movement execution tasks, but it remains unclear how these neural responses relate to distinct reward and sensory-error based motor learning mechanisms (Krigolson et al. 2008; Torrecillos et al. 2014; MacLean et al. 2015). Here we identified neural signatures of sensory error and reward feedback processing in motor learning using separate learning paradigms that produce comparable changes in behavior.

In theories of motor adaptation, SPE occurs when the sensory consequences of motor commands differ from the predicted outcomes. SPE is thought to occur in visuomotor rotation (VMR) paradigms in which visual feedback of hand position is rotated relative to the actual angle of reach. Adaptation, in which motor output is adjusted to compensate for perturbations, is thought to be driven largely by SPE in these tasks (Izawa and Shadmehr 2011; Marko et al. 2012). Sensory error feedback activates brain regions including primary sensory motor areas, posterior parietal cortex, and cerebellum (Inoue et al. 2000,2016; Krakauer et al. 2004; Diedrichsen et al. 2005; Bédard and Sanes 2014). Tanaka et al. (2009) propose that SPEs computed by the cerebellum produce adaptation via changes in synaptic weighting between the posterior parietal cortex and motor cortex. Furthermore, strategic aiming also contributes to behavioural compensation for visuomotor rotations in a manner that is largely independent from the automatic visuomotor recalibration that is driven by cerebellar circuits (Mazzoni and Krakauer 2006; Benson et al. 2011; Taylor et al. 2014; McDougle et al. 2016).

Recent research suggests that a reward based learning process can also contribute to motor adaptation in parallel to a sensory-error based learning system, and that reward feedback can drive motor learning even in the absence of sensory error feedback (Izawa and Shadmehr 2011; Therrien et al. 2015; Holland et al. 2018). Reward learning has been isolated experimentally by providing participants with only binary reward feedback, indicating success or failure, without visual feedback of hand position. Reward-based motor learning has been modelled as reinforcement learning, which maps actions to abstract representations of reward or success, rather than to the sensory consequences of action. In reinforcement learning theory, if the outcome of an action is better than the predicted outcome, a positive reward prediction error (RPE) occurs which drives an increase in the agent’s estimate of expected value of that reward outcome, along with an increase in the future likelihood of selecting that particular action. Conversely, if the outcome is worse than expected, a negative RPE diminishes the estimated value and likelihood of selecting that action. Phasic dopaminergic signaling in the VTA and striatum is consistent with encoding of RPE (Glimcher 2011), and dopaminergic activity has been implicated in reward based motor learning (Galea et al. 2013; Pekny et al. 2015).

EEG has been used to identify neural correlates of error monitoring, but few studies have employed motor adaptation tasks. An event-related potential (ERP) known as the feedback related negativity (FRN) has been proposed to reflect processing of RPE in the context of reinforcement learning. There is evidence that FRN reflects activity in anterior cingulate or supplementary motor cortical areas, and the reinforcement learning theory of the FRN states that it is driven by phasic dopamin ergic signaling of RPE (Holroyd and Coles 2002; Nieuwenhuis et al. 2004; Walsh and Anderson 2012; Heydari and Holroyd 2016). Other accounts attribute the FRN to processes involving conflict monitoring or general prediction error as opposed to RPE (Alexander and Brown 2011; Baker and Holroyd 2011a; Botvinick 2011). Motor adaptation paradigms are particularly opportune to test the reinforcement learning theory of the FRN as they afford dissociation between reinforcement learning processes and learning through sensory prediction error.

The FRN potential is superimposed on the P300, a well characterized positive ERP component that peaks later than FRN, and with a more posterior scalp distribution. It has been proposed that the P300 reflects the updating of a model of stimulus context (Donchin and Coles 1988). Both the FRN and the P300 have been observed in response to errors in motor tasks (Krigolson et al. 2008; Torrecillos et al. 2014; MacLean et al. 2015; Reuter et al. 2018; Savoie et al. 2018). In this paper we describe experiments in which we isolated and compared EEG responses to both reward and sensory error feedback using separate adaptation paradigms that produced comparable changes in behavior. We tested the idea that the FRN is a neural signature of feedback processing that specifically supports reward based motor learning, while the P300 reflects a process which is generally related to feedback processing but is particularly important for sensory error based learning.

## Material and Methods

### Experimental Design and Statistical Analysis

Participants made reaching movements toward a visual target while holding the handle of a robotic arm. A screen displayed visual information related to the task but occluded vision of the arm. The setup and procedure regarding the reaching movements is described under “Apparatus/Behavioral Task”. Feedback pertaining to reach angle was provided at movement endpoint, and feedback was manipulated such that participants adapted their reach direction to compensate for the manipulations. Participants were instructed that each reach terminating within the target would be rewarded with a small monetary bonus. Participants underwent alternating experimental blocks of a reward learning task and a visuomotor rotation task. The design for each task is described under “reward learning task” and “visuomotor rotation task” methods subsections, respectively.

In the visuomotor rotation task, a cursor appeared at movement endpoint to represent the position of the hand. In a randomly selected 50% of trials, a perturbation was imposed, such that the cursor feedback was rotated around the starting position by a fixed angle, indicating a reach angle that was shifted relative to the unperturbed feedback. The magnitude of the perturbation was varied across blocks, and was either .75 deg or 1.5 deg. Behaviourally, we tested for trial by trial adaptive responses that compensated for the perturbations. We compared the neural responses to rotated and non-rotated feedback to assess the neural correlates of processing sensory error feedback during adaptation, and we tested whether this effect was modulated by the size of the perturbations. The perturbations were small relative to the size of the target, such that participants nearly always landed in the target, fulfilling the goal of the task and earning a monetary reward, even on the perturbed trials. Thus, reward and task error were constant between perturbed and non-perturbed feedback, and by comparing the two conditions we could assess the neural and behavioural response to sensory error without the confounds of reward or task error processing.

In the reward learning task, no cursor appeared to indicate the position of the hand, but instead binary feedback indicated whether or not participants succeeded in hitting the target and earning monetary reward. This allowed us to assess the neural and behavioral responses to reward feedback in isolation from sensory error processing, as visual information revealing the position of the hand was not provided. Reward was delivered probabilistically, with a higher probability of reward for reaches in one direction than the other, relative to participants’ recent history of reach direction. This was intended to induce adaptation such that participants would adjust their reaches towards the direction that was rewarded at a higher probability. The overall reward frequency was controlled and manipulated across blocks, so that participants experienced both reward and non-reward feedback in both low and high overall reward frequency conditions. We compared the neural responses to reward and non-reward feedback to assess the neural correlates of reward processing during adaptation. We compared the responses to frequent and infrequent feedback to assess effects related to expectation, under the assumption that outcomes which occurred less frequently in a given block would violate expectations more strongly (Reward feedback in the high reward frequency condition and non-reward feedback in the low reward frequency conditions were deemed “frequent”, while non-reward feedback in the high reward frequency condition and reward feedback in the low reward frequency condition were deemed “infrequent”).

The trial averaging procedures used to estimate the neural responses to various feedback conditions are described under “Event Related Component Averaging”, and the analysis of these neural responses is described under “P300 Analysis” and “Feedback Related Negativity Analysis”.

Results of statistical tests are reported in the Results section, under “Behavioral Results”, “Feedback Related Negativity Results”, and “P300 Results”. Linear relationships between behavioural and EEG measures were assessed using robust regression, implemented by the Matlab fitlm function with robust fitting option. This method uses iteratively reweighted least squares regression, assigning lower weight to outlier data points. Student’s t-tests were performed using MATLAB R2016b, and Lilliefors test was used to test the assumption of normality. In the case of non-normal data, Wilcoxin signed rank test was used to test pairwise differences. Repeated measures analyses of variance (ANOVAs) were conducted using IBM SPSS Statistics version 25. For all ANOVAs, Mauchly’s test was used to validate the assumption of sphericity.

### Participants

A total of n=20 healthy, right-handed participants were included in our study (23.21 ± 3.09 years old, 12 females). Three participants underwent the experimental procedure but were excluded due to malfunction of the EEG recording equipment. One participant who reported that they performed movements based on a complex strategy that was unrelated to the experimental task was excluded. Participants provided written informed consent to experimental procedures approved by the Research Ethics Board at The University of Western Ontario.

### Experimental Procedure

Participants first performed a block of 50 practice trials. After the practice block the experimenter fitted the EEG cap and applied conductive gel to the electrodes before beginning the behavioral task (see “EEG Data Acquisition” below). The behavioral procudure consisted of 8 experimental blocks, including four 115-trial blocks of a reward learning task and four 125-trial blocks of a visuomotor rotation task. The order of the blocks alternated between the two task types but was otherwise randomized. Participants took self-paced rests between blocks.

### Apparatus/Behavioral Task

Participants produced reaching movements with their right arm in a horizontal plane at chest height while holding the handle of a robotic arm (InMotion2, Interactive Motion Technologies, Massachusetts, United States; Fig 1). Position of the robot handle was sampled at 600 Hz. A semi-silvered mirror obscured vision of the arm and displayed visual information related to the task. An air-sled supported each participant’s right arm.

**Figure 1:**
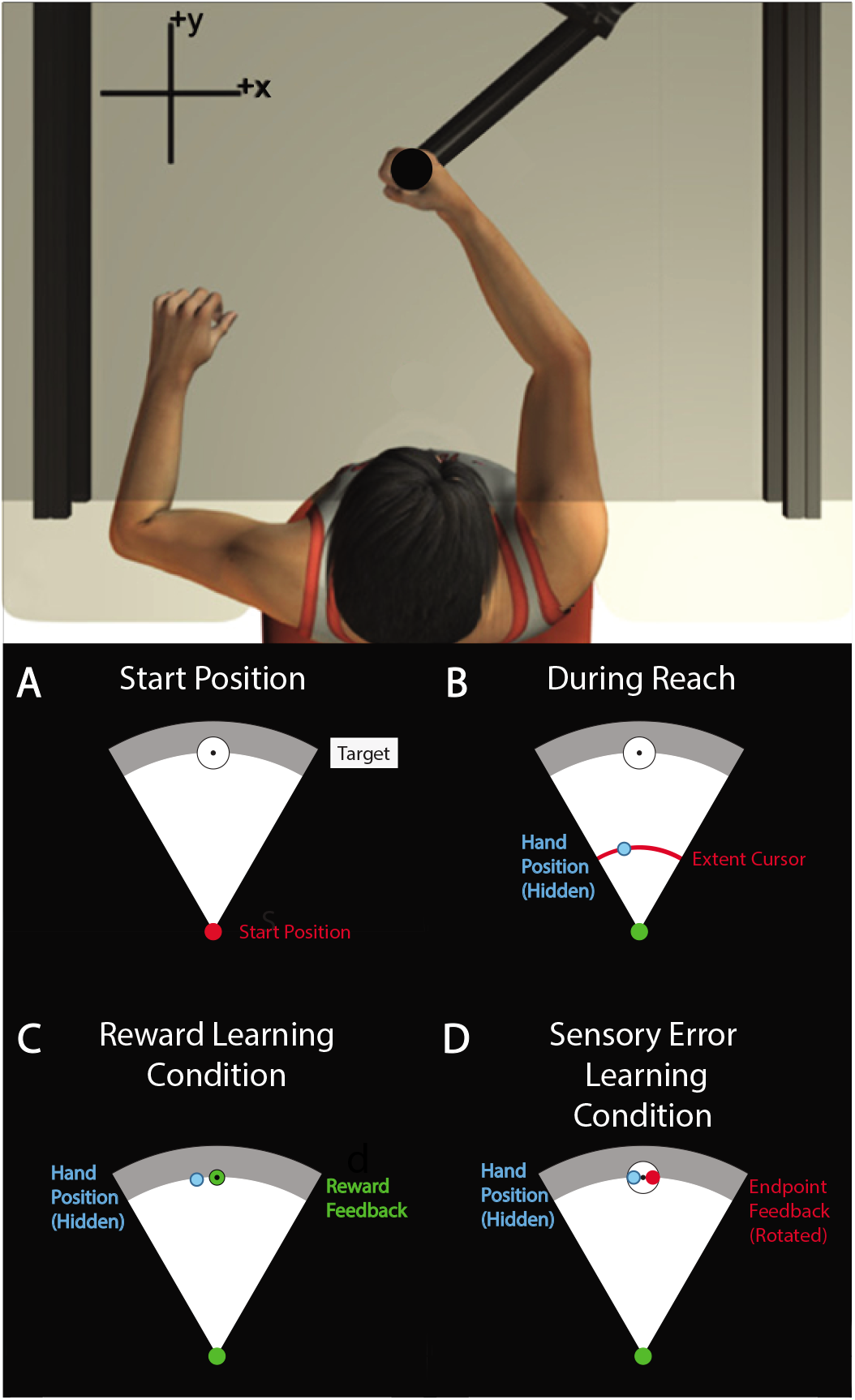
Experimental setup. A, Participants reached to visual targets while holding the handle of a robotic arm. Vision of the arm was obscured by a screen that displayed visual information related to the task. B, During reaches, hand position was hidden but an arc shaped cursor indicated the extent of the reach without revealing reach angle. Feedback was provided at reach endpoint. C, In the reward learning condition, binary feedback represented whether reaches were successful or unsuccessful in hitting the target by turning green or red, respectively. Reach adaptation was induced by providing reward for movements that did not necessarily correspond to the visual target. D, In the visuomotor rotation condition, feedback represented the endpoint position of the hand. Adaptation was induced by rotating the angle of the feedback relative to the actual reach angle.

Participants began each trial with their hand at a start position in front of their chest at body midline (mid-saggital plane). The start position was displayed using a red circle with a diameter of 1 cm (Fig 1a). A white circular target was displayed 14 cm away from the start position (Fig 1a). A cursor indicated hand position only while the hand was within the start circle. The start position turned green to cue the onset of each reach once the handle had remained inside the start position continuously for 750 ms. Participants were instructed that they must wait for the cue to begin each reach, but that it was not necessary to reach immediately or react quickly upon seeing the cue.

Participants were instructed to make forward reaches and to stop their hand within the target. An arc shaped cursor indicated reach extent throughout the movement, without revealing the angle of the hand relative to the start position. In the first 5 baseline trials of each block, continuous position feedback was provided and consisted of an additional circular cursor indicated the position of the hand throughout the reach. In all subsequent reaches for each block, the cursor indicating hand position disappeared when the hand left the start position, and only the arc shaped cursor indicating movement extent was shown. A viscous force field assisted participants in braking their hand when the reach extent was greater than 14 cm.

The robot ended each movement by fixing the handle position when the hand velocity decreased below 0.03 m/s. During this time, while the hand was fixed in place (for 700 ms) visual feedback of reach angle was provided. Feedback indicated either reach endpoint position, a binary reward outcome, or feedback of movement speed (see below). Visual feedback was then removed and the robot guided the hand back to the start position. During this time no visual information relating to hand position was displayed.

Reach endpoint was defined as the position at which the reach path intersected the perimeter of a circle (14 cm radius), centered at the start position. Reach angle was calculated as the angle between a line drawn from the start position to reach endpoint and a line drawn from the start position to the center of the target, such that reaching straight ahead corresponds to 0 deg and counter-clockwise reach angles are positive (Fig 1a). Feedback about reach angle was provided either in the form of endpoint position feedback or binary reward feedback. The type of feedback, as well as various feedback manipulations, varied according to the assigned experimental block type (See “Reward Learning Condition” and “Visuomotor Rotation Condition”). Endpoint position feedback consisted of a stationary cursor indicating the position of movement endpoint, while reward feedback consisted of the target turning either red or green to indicate that the reach endpoint missed or hit the target, respectively.

Movement duration was defined as the time elapsed between the hand leaving the start position and the moment hand velocity dropped below 0.03 m/s. If movement duration was greater than 700 ms or less than 450 ms no feedback pertaining to movement angle was provided. Instead, the gray arc behind the target turned blue or yellow to indicate that the reach was too slow or too fast, respectively. Participants were informed that movements with an incorrect speed would be repeated but would not otherwise affect the experiment.

To minimize the impact of eye-blink related EEG artifacts, participants were asked to fixate their gaze on a black circular target in the center of the reach target and to refrain from blinking throughout each arm movement and subsequent presentation of feedback.

### Practice Block

Each participant completed a block of practice trials before undergoing the reward learning and visuomotor rotation experiments. Practice continued until participants achieved 50 movements within the desired range of movement duration. Continuous position feedback was provided during the first five trials, and only endpoint position feedback was provided for the subsequent 10 trials. After these initial 15 trials, no position feedback was provided outside the start position.

### Reward Learning Task

Binary reward feedback was provided to induce adaptation of reach angle. Each participant completed 4 blocks in the reward learning condition. We manipulated feedback with direction of intended learning and reward frequency as factors using a 2×2 design (direction of learning x reward frequency) across blocks. For each direction of intended learning (clockwise and counter-clockwise), each participant experienced a block with high reward frequency and a block with low reward frequency. Participants performed blocks with each direction of intended learning to control for bias in reach direction. Reward frequency was manipulated to assess effects related to expectation, under the assumption that outcomes which occurred less frequently in a given block would violate expectations more strongly. Each reward learning block continued until the participant completed 115 reaches with acceptable movement duration. Participants reached towards a circular target 1.2 cm (4.9 deg) in diameter. The first 15 reaches were baseline trials during which the participant did not receive reward feedback. Continuous position feedback was provided during the first five trials, and only endpoint position feedback was provided for the subsequent 10 trials. After these baseline trials, no position feedback was provided, and binary reward feedback was provided at the end of the movement. Participants were told that they would earn additional monetary compensation for reaches that ended within the target, up to a maximum of CAD$10 for the whole experiment. Participants were told that rewarded and unrewarded reaches would be indicated by the target turning green and red, respectively.

Unbeknownst to participants, reward feedback was delivered probabilistically. The likelihood of reward depended on the difference between the current reach angle and the median reach angle of the previous 10 reaches. In the high reward frequency condition, reward was delivered with probability of 100% if the difference between the current reach angle and the running median was in the direction of intended learning, and at a probability of 30% otherwise (eq. 1). When the running median was at least 6 deg away from zero in the direction of intended learning, reward was delivered at a fixed probability of 65%. This was intended to minimize conscious awareness of the manipulation by limiting adaptation to ± 6 deg. In the low reward frequency condition, reward was similarly delivered at a probability of either 70% or 0% (eq. 2). When the running median was at least 6 deg away from zero in the direction of intended learning, reward was delivered at a fixed probability of 35%. Reach angle and feedback throughout a representative experimental block is shown in Figure 2.

**Figure 2:**
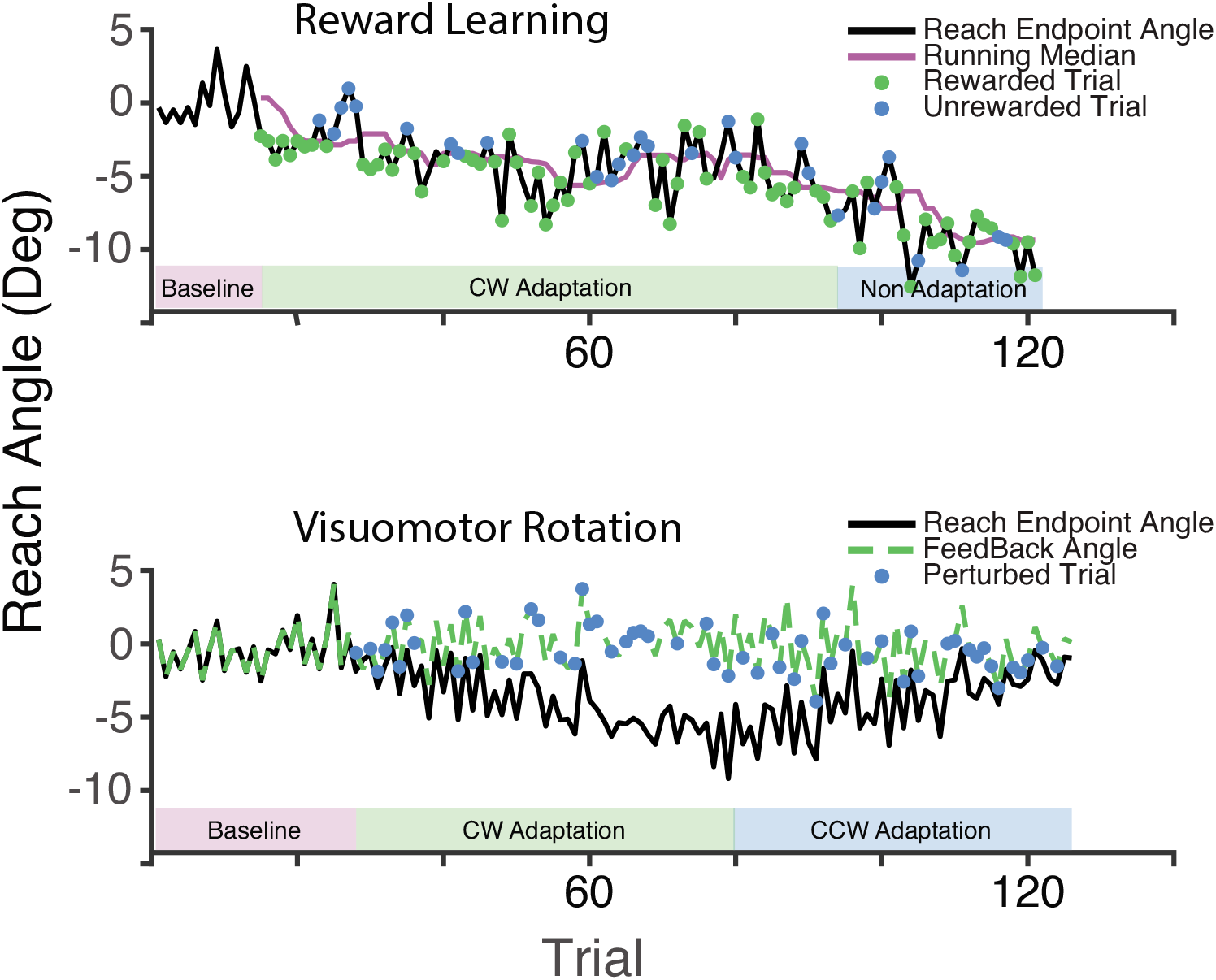
Reach angles of a representative participant. A) We show the reward learning block assigned to the clockwise adaptation with high reward frequency condition. Reaches were rewarded with 100.0% probability for reach angles less than the median of the previous 10 reaches, and with 30.0% probability for reach angles greater than this running median. Reward was delivered at a fixed probability of 65.0% when the running median was less than −6 degs, indicated by the ‘Non-Adaptation’ portion of the block. B) The visuomotor rotation block assigned to the 1.5 degree rotation condition is shown. The rotation is imposed randomly in 50% of trials. The rotation is initially counterclockwise but reverses when the mean of the previous five reach angles becomes less than −6.0 deg.

We employed this adaptive, closed loop reward schedule so that the overall frequency of reward was controlled. While participants adapted their reach angle to the task, the task adapted to the changing reach angle, as each reach was assessed relative to the recent history of reaches. This allowed us to assess correlations between neural measures and behavior without confounding learning and reward frequency.

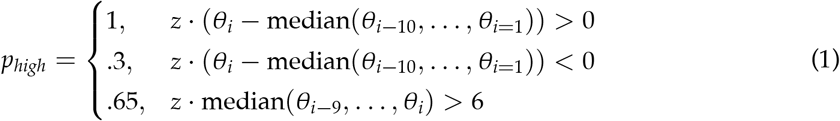

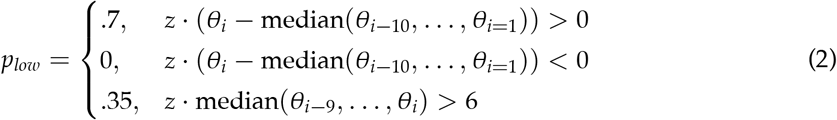

Where *p* is probability of reward described separately for the high and low reward frequency conditions, *θ* is the reach angle on trial *i, z* = 1 for counter-clockwise learning blocks, and *z* = −1 for clockwise learning blocks.

### Visuomotor Rotation Task

Endpoint feedback was rotated relative to the actual reach angle to induce sensory-error based adaptation. Each participant completed 4 blocks in the visuomotor rotation condition. We manipulated feedback with initial rotation direction and perturbation size as factors using a 2×2 design across blocks. For each direction of initial rotation (clockwise and counter-clockwise) each participant experienced a block with large rotation (1.5 deg) and a block with small rotation (.75 deg). Each block continued until participants completed 125 reaches within acceptable movement duration limits. Participants reached towards a circular target 2.5 cm (10.2 deg) in diameter. Participants first performed 25 baseline reaches during which position feedback reflected veridical reach angle. A cursor continuously indicated hand position during the first 10 trials. A static cursor indicated movement endpoint position for the subsequent 15 trials. After the baseline reaches, the adaptation portion of each block began unannounced to participants and continued until the participant performed 100 additional reaches of acceptable movement duration.

During the adaptation trials, endpoint position feedback was provided that did not necessarily correspond to the true hand position. Participants were instructed that endpoint feedback within the target would earn them bonus compensation, but no explicit reward feedback was provided during the experiment. To determine the feedback angle in the small and large perturbation conditions, in a randomly selected 50% of trials we added a rotation of 0.75 deg or 1.5 deg, respectively, to the true reach angle. In addition, on every trial, we subtracted an estimate of the current state of reach adaptation (eq. 3).

To illustrate the purpose of subtracting a running estimate of the current state of adaptation, we can consider the case in which reach angle was adapted by +2 deg through cumulative exposure to a -.75 deg rotation. If the state of adaptation is accurately estimated and subtracted from the true reach angle, then a reach angle of +2 deg will result in either unperturbed feedback at 0 deg or rotated feedback at -.75 deg. The online estimate of adaptation consisted of a running average of the previous 6 reach angles and a model of reach adaptation which assumed that participants would adapt to a fixed proportion of the reach errors experienced during the previous 3 trials. A windowed average centered around the current reach angle could serve as a low pass filter to estimate the current state of reach adaptation, but the running average during the experiment was necessarily centered behind the current reach angle. Thus, an online model was necessary to account for adaptation that would occur in response to errors experienced on trials included in the running average. Mis-estimation of the learning rate in the online model would lead to systematic bias in the feedback angle, while a perfect model would lead to unperturbed feedback that is distributed around 0 deg with variance that reflects the natural movement variability. An adaptation rate of 0.25 was chosen for the online model on the basis of pilot data.

This design allowed us to compare perturbed and unperturbed feedback in randomly intermixed trials, which is generally advantageous in ERP experiments. Previous studies have imposed a fixed perturbation throughout a block of trials and compared early trials, in which the visual error is large, to late trials, in which the error has been minimized through adaptation (Tan et al. 2014; MacLean et al. 2015). However, in such designs the independent variable is not experimentally controlled, and differences in neural response might be attributed to changes in the state of adaptation or simply habituation to feedback, as opposed to sensory error per se. Another alternative is to impose random rotations in either direction, but previous work has demonstrated that neural and behavioural responses are larger for consistent perturbations, presumably because the sensorimotor system attributes variability in feedback to noise processes and downweights adaptive responses accordingly (Tan et al. 2014).

As in the reward learning condition, we sought to limit the magnitude of adaptation to 6 deg in an attempt to minimize conscious awareness of the manipulation. This limit was implemented by reversing the direction of the perturbation whenever the average reach angle in the previous six movements differed from zero by at least 6 deg in the direction of intended reach adaptation. Reversing the direction of the perturbation caused participants to adapt in the opposite direction. Reach angle and feedback angle throughout a representative experimental block is shown in Figure 2.

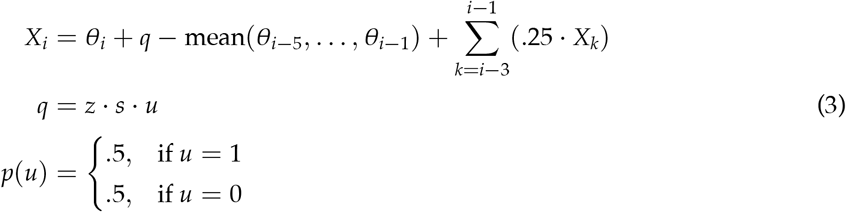

*X* denotes feedback angle, *θ* denotes reach angle, and *q* denotes the perturbation. *z* denotes the direction of the perturbation (*z* = 1 for counter-clockwise perturbations, and z = −1 for clockwise perturbations). *s* denotes the size of the perturbation (.75 deg or 1.5 deg in the small and large error conditions, respectively). *u* is a discrete random variable that is realized as either 1 or 0 with equal probability (50%).

### EEG Data Acquisition

EEG data were acquired from 16 cap-mounted electrodes using an active electrode system (g.Gamma; g.tec Medical Engineering) and amplifier (g.USBamp; g.tec Medical Engineering). We recorded from electrodes placed according to the 10-20 system at sites Fz, FCz, Cz, CPz, Pz, POz, FP1, FP2, FT9, FT10, FC1, FC2, F3, F4, F7, F8, referenced to an electrode placed on participants’ left earlobe. Impedances were maintained below 5 kU. Data were sampled at 4800 Hz and filtered online with band-pass (0.1-1,000 Hz) and notch (60 Hz) filters. The amplifier also recorded data from a photo-diode attached to the display monitor to determine the timing of stimulus onset.

### Behavioral Data Analysis

#### Reward learning task

Motor learning scores were calculated for each participant as the difference between the average reach angle in the counter-clockwise learning blocks and the average reach angle in the clockwise learning blocks. We chose to assess reach angle throughout the entire task, as opposed to only in a window of trials at the end of each block, primarily because reach direction was often unstable and a smaller window was susceptible to drift. Furthermore, this metric of learning measured not only the final state of adaptation but also reflected the rate of adaptation throughout the block without assuming a particular function for the time course of learning. Lastly, this metric was not dependent on the choice of a particular subset of trials.

We excluded baseline trials and trials that did not meet the movement duration criteria, as no feedback related to reach angle was provided on these trials (6.5% of trials in the VMR task, 7.4% of trials in the RL task.)

#### Visuomotor rotation task

To quantify trial-by-trial learning we first calculated the change in reach angle between successive trials, as in eq. 4.

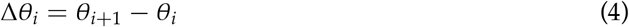

We then performed a linear regression on Δ*θ_i_* with the rotation imposed on the trial *i* as the predictor variable. The rotation was either 0, ± 0.75, or ± 1.5 deg. This regression was performed on an individual participant basis, separately for each of the four visuomotor rotation conditions (corresponding to feedback rotations of −1.5, −0.75, 0.75, and 1.5 deg). For these regressions, we excluded trials that did not meet the duration criteria or that resulted in a visual error of greater than 10 deg (M=2.65 trials per participant, SD=4.3), as these large errors were thought to reflect execution errors or otherwise atypical movements. We took the average of the resulting slope estimates across blocks, multiplied by −1, as a metric of learning rate for each participant, as it reflects the portion of visual errors that participants corrected with a trial-by-trial adaptive process. Based on simulations of our experimental design using a standard memory updating model (Thoroughman and Shadmehr 2000) (not described here), we found that it was necessary to perform the regression separately for each rotation condition, as collapsing across the different rotation sizes and directions could introduce bias to the estimate of learning rate.

### EEG Data Denoising

EEG data were resampled to 480 Hz and filtered offline between 0.1-35 Hz using a second order Butterworth filter. Continuous data were segmented into 2 second epochs time-locked to feedback stimulus onset at 0 ms (time range: −500 to +1500 ms). Epochs containing artifacts were removed using a semi-automatic algorithm for artifact rejection in the EEGLAB toolbox (see Delorme and Makeig (2004) for details). Epochs flagged for containing artifacts as well as any channels with bad recordings were removed after visual inspection. Subsequently, extended infomax independent component analysis was performed on each participant’s data (Delorme and Makeig 2004). Components reflecting eye movements and blink artifacts were identified by visual inspection and subtracted by projection of the remaining components back to the voltage time series.

### Event Related Component Averaging

We computed event related potentials (ERPs) on an individual participant basis by trial averaging EEG time series epochs after artifact removal. We selected trials corresponding to various feedback conditions in each task as described in the following sections. For each feedback condition ERPs were computed on an individual participant basis separately for recordings from channels FCz and Pz. All ERPs were baseline corrected by subtracting the average voltage in the 75 ms period immediately following stimulus onset. We chose to use a baseline period following, as opposed to preceding, stimulus onset because stimuli were presented immediately upon movement termination, and the period before stimulus presentation was more likely to be affected by movement related artifacts. Importantly, we did not observe any ERPs with onsets within the baseline period. Trials in which reaches did not meet the movement duration criteria were excluded, as feedback relevant to reach adaptation was not provided on these trials (6.5% of trials in the VMR task, 7.4% OF trials in the RL task.).

#### Reward Learning Task

ERPs from the reward learning condition were used to examine the neural correlates of reward feedback processing. We computed ERPs separately for feedback conditions corresponding to “frequent reward”, “infrequent reward”, “frequent non-reward”, and “infrequent non-reward”. Expectancy was determined on the basis of the reward frequency condition: reward in the high reward frequency and non-reward in the low reward frequency conditions were deemed frequent, while reward in the low reward frequency and non-reward in the high reward frequency conditions were deemed infrequent (Holroyd and Krigolson 2007).

#### Visuomotor Rotation Task

ERPs elicited by visuomotor rotation were used to examine the neural correlates of sensory error feedback processing. We created trial averaged ERP responses for trials with rotated feedback and trials with non-rotated feedback separately for the 0.75 deg and 1.5 deg rotation conditions. Size of rotation was varied across blocks, while within each block rotations were imposed in a randomly selected 50% of trials. The resulting ERPs are identified by the conditions “rotated 0.75 deg”, “non-rotated 0.75 deg”, “rotated 1.5 degs” and “non-rotated 1.5 deg”.

In order to test for effects of absolute endpoint error, which is determined not only by visuomotor rotation but also movement execution errors, we sorted trials in the adaptation portion of the visuomotor rotation blocks by the absolute value of the angle of visual feedback relative to the center of the target. We created “most accurate” ERPs for each participant by selecting the 75 trials with the smallest absolute feedback angle, and “least accurate” ERPs by selecting the 75 trials with the largest absolute feedback angle.

We computed ERPs to test a correlation, across participants, between behavioral learning rate and the average neural response to feedback during adaptation to visuomotor rotation. These ERPs, labelled as belonging to the “adaptation” condition, included all trials in the “rotated 0.75 deg”, “non-rotated 0.75 deg”, “rotated 1.5 degs” and “non-rotated 1.5 deg” conditions.

### Feedback Related Negativity Analysis

The FRN was analyzed using a difference wave approach with event related potentials (ERPs) recorded from FCz, where it is typically largest (Miltner et al. 1997; Holroyd and Krigolson 2007; Pfabigan et al. 2011). By computing a difference wave between non reward and reward outcomes, it is possible to capture a bidirectional voltage response to reward vs non reward feedback. Although the feedback related negativity is classically characterized by a negative voltage peak following non reward feedback, multiple lines of evidence suggest that voltage increase in response to reward outcomes also contributes to the variance captured by the difference wave approach, despite not producing a distinct peak (Baker and Holroyd 2011b; Carlson et al. 2011; Walsh and Anderson 2012; Becker et al. 2014; Heydari and Holroyd 2016). Furthermore, difference waves can be computed separately for frequent and infrequent outcomes, capturing effects related to reward prediction error while removing effects of pure surprise through subtraction. Difference waves were computed for each participant by subtracting ERPs corresponding to unsuccessful outcomes from those corresponding to successful outcomes. FRN amplitude was determined as the mean value of the difference wave between 200-350 ms after feedback presentation. This time window was chosen a priori on the basis of previous reports (Walsh and Anderson 2012). To test for the statistically significant presence of the FRN for each difference wave, we submitted the mean value between 200-350 ms after feedback presentation to a t-test against zero.

#### Visuomotor Rotation Task

First, we created difference waves to test whether the rotations imposed on randomly selected trials elicited FRN. The “rotated 0.75 deg” event related potentials (ERPs) were subtracted from the “non-rotated 0.75 deg” ERPs to create a “small VMR” difference wave. The “rotated 1.5 deg” ERPs were subtracted from the “non-rotated 1.5 deg” ERPs to create a “large VMR” difference wave.

Next, we created a difference wave to test whether a FRN was observable by comparing trials where the endpoint feedback was furthest from the center of the target from those where feedback was closest to the center of the target. The “least accurate” ERPs were subtracted from the “most accurate” ERPs to create an “endpoint error” difference wave. We hypothesized that sensory error feedback would not elicit a FRN, and as such we tested for the FRN using this second approach to more thoroughly confirm our hypothesis that the FRN would not be observed in the visuomotor rotation task.

#### Reward Learning Task

The frequent non-reward event related potential (ERP) was subtracted from the frequent reward EPR to create a “frequent” difference wave, and the infrequent non-reward ERP was subtracted from the infrequent reward EPR to create an “infrequent” difference wave.

We used a t-test to test the difference between FRN amplitude for the “frequent” and “infrequent” difference waves.

### P300 analysis

To analyze the P300 we used event related potentials (ERPs) recorded from channel Pz, where it is typically largest (Fabiani et al. 1987; Hajcak et al. 2005; Polich 2007; MacLean et al. 2015). We calculated P300 amplitude using base-to-peak voltage difference. The temporal ROIs for the peak and base were determined using grand averages computed across participants and conditions for each task (see “Visuomotor Rotation task”, and “Reward Learning Task”, below). P300 peak was defined as the maximum peak occurring 250-500 ms after stimulus onset, which always corresponded to the largest peak in the analyzed epoch. P300 base was defined as the minimum preceding peak that occurred at least 100 ms after stimulus onset. For each subject, peak and base voltages were calculated separately for each condition ERP as the average voltage within 50 ms windows centered around the temporal ROIs defined at the group level. P300 amplitude was then determined as the difference between peak and base voltage.

#### Visuomotor Rotation task

P300 amplitude was calculated in four conditions using the “rotated 0.75 deg”, “non-rotated 0.75 deg”, “rotated 1.5 deg” and “non-rotated 1.5 deg”, event related potentials (ERPs). Temporal ROIs were determined, as described above, by aggregating all trials across participants and the four conditions into a single set and averaging to produce an “aggregate grand average from trials” waveform. This approach allows for data driven ROI selection without inflated Type-I error rate, and has been shown to be insensitive to trial number asymmetry across conditions (Brooks et al. 2017). We tested for differences in P300 amplitude related to visuomotor rotation using two-way repeated measures ANOVA with factors rotation (levels: non-rotated, rotated) and rotation magnitude (levels: 0.75 deg, 1.5 deg).

#### Reward Learning Task

P300 amplitude was calculated in four conditions using the “infrequent reward”, “frequent reward”, “infrequent non-reward”, and “frequent non-reward” feedback condition event related potentials (ERPs), described above. We found that the waveform morphology was considerably different for the ERPs elicited by reward feedback and those elicited by non-reward feedback, in part due to the feedback related negativity. As such we defined temporal ROIs separately for the reward conditions (infrequent reward, frequent reward) and the non-reward conditions (“infrequent non-reward”, and “frequent non-reward”). In both cases, temporal ROIs were determined, as described above, by aggregating all trials across participants and the corresponding two conditions into a single set and averaging to produce an “aggregate grand average from trials” waveform. This ROI selection method has only been shown to be necessarily unbiased when all conditions display similar waveform morphology and are grouped together (Brooks et al. 2017). For this reason, we repeated our analysis using a common method of selecting peaks for each individual participant and condition ERP after matching the number of trials across conditions, which produced similar results.

We tested for differences in P300 amplitude between feedback conditions using 2 way repeated measures ANOVA with factors reward (levels: rewarded, non-rewarded) and expectancy (levels: infrequent, frequent).

## Results

### Behavioral Results

#### Reward Learning Task

In the reward learning task participants adapted their reach angle on the basis of binary reward feedback (Fig 3). We calculated a reward learning score for each subject by subtracting the average reach angle in the clockwise learning condition from that in the counter-clockwise learning condition, excluding the baseline trials, such that the average reward learning score would be approximately zero if participants did not respond to the reward feedback in any way. We observed a mean reward learning score of 5.47 (SD=4.66), which is significantly greater than zero (one-sample t-test; t(19)=5.25, p<.001). Participants received reward on 67.0% (SD=4.9%) of trials in the high frequency condition and 38.6% (SD=4.3%) of trials in the low frequency condition.

**Figure 3:**
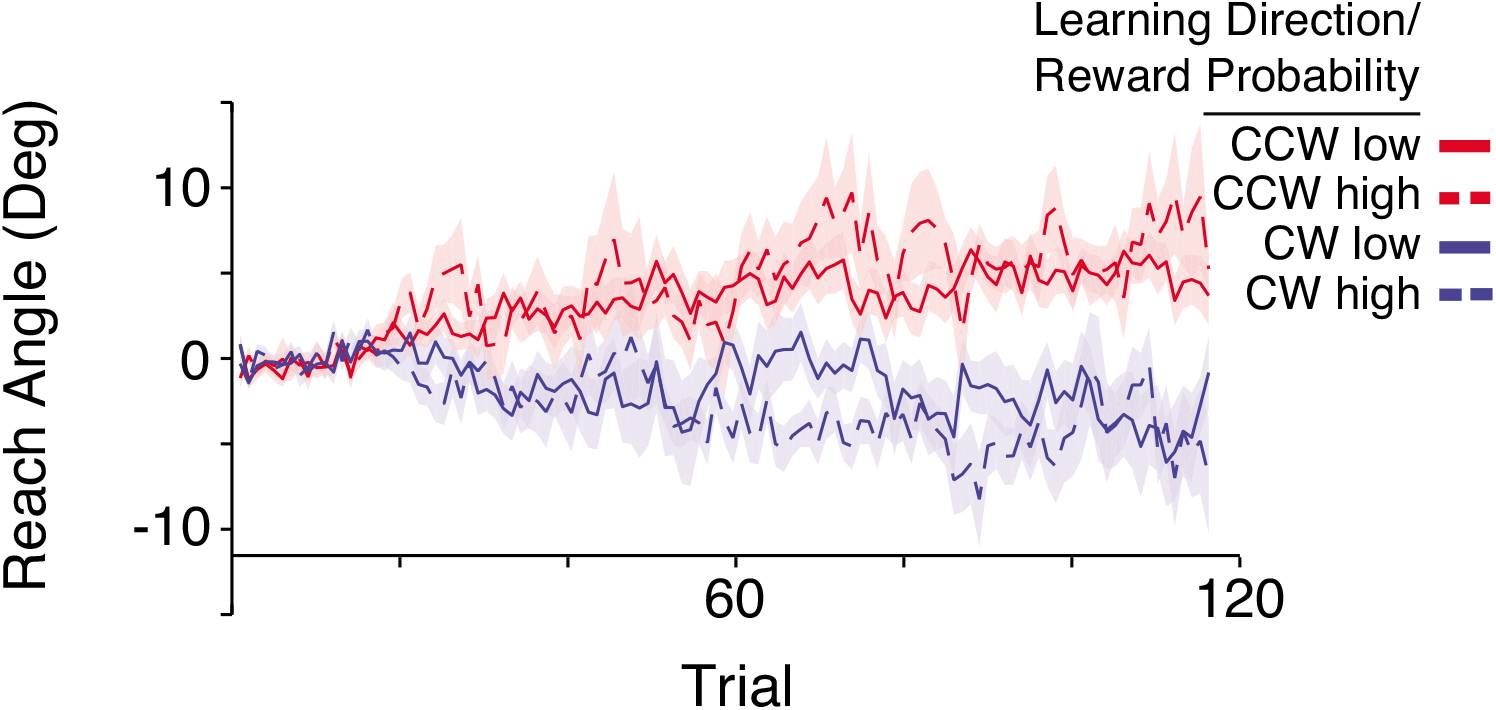
Participants adapted their reach angle in the reward learning condition. Group average reach angles in the reward learning conditions are plotted. Each participant completed four blocks. For each direction of intended learning (clockwise and counter clockwise), each participant completed a block in the high reward frequency (65%) condition and a block in the low reward frequency (35%) condition. Shaded regions are ± 1 SEM.

#### Visuomotor rotation task

In the visuomotor rotation task participants received endpoint cursor feedback and adapted their reach angles in response to the rotated cursor feedback imposed on randomly selected trials. To estimate trial-by-trial learning rates for individual participants we quantified the linear relationship between the change in reach angle after each trial with the rotation imposed on the preceding trial as the predictor variable, separately for each rotation condition (−1.5, −0.75, 0.75, and 1.5 deg). We took the average of the resulting slope estimates and multiplied it by negative one to obtain a measure of learning rate. This metric reflects the proportion of visuomotor rotation that each participant corrected with a trial-by-trial adaptive process. The mean learning rate was 0.49 (SD=0.46), which was significantly different from zero (one-sample t-test; t(19)=4.8, p<.001). This indicates that participants corrected for visual errors on a trial-by-trial basis. Figure 4 shows the average change in reach angle for each size and direction of the imposed cursor rotation.

**Figure 4:**
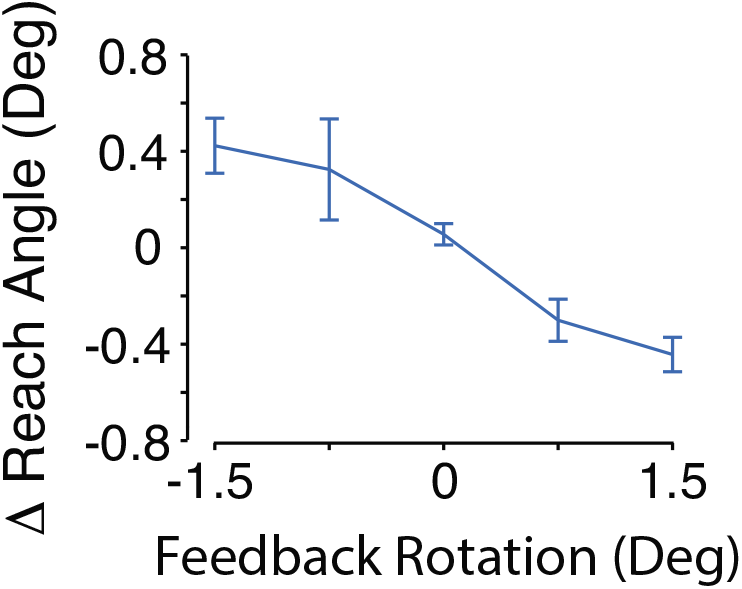
Participants adapted their reach angle on a trial-by-trial basis in the visuomotor rotation condition. The average change in reach angle between subsequent pairs of trials is plotted for each size and direction of rotation imposed on the preceding trial. The average change in reach angle is in all cases opposite to the rotation, indicating that participants adapted their reaches to counteract the perturbations.

### Feedback Related Negativity Results

#### Reward learning task

Figure 5a shows the event related potentials (ERPs) recorded from electrode FCz during the reward learning condition, averaged across participants. The mean value of the “frequent” difference wave recorded from FCz between 200-350 ms was significantly different from zero (M=5.34 *μ*V, SD=4.11, t(19)=5.81, p<.001, one-sample t-test), indicating that frequent feedback elicited a FRN in our reward learning task. The mean value of the “infrequent” difference wave was also significantly larger than zero (M=7.09 *μ*V, SD=2.76, t(19)=11.47, p<.001, one-sample t-test), indicating that infrequent feedback also elicited a FRN.

**Figure 5:**
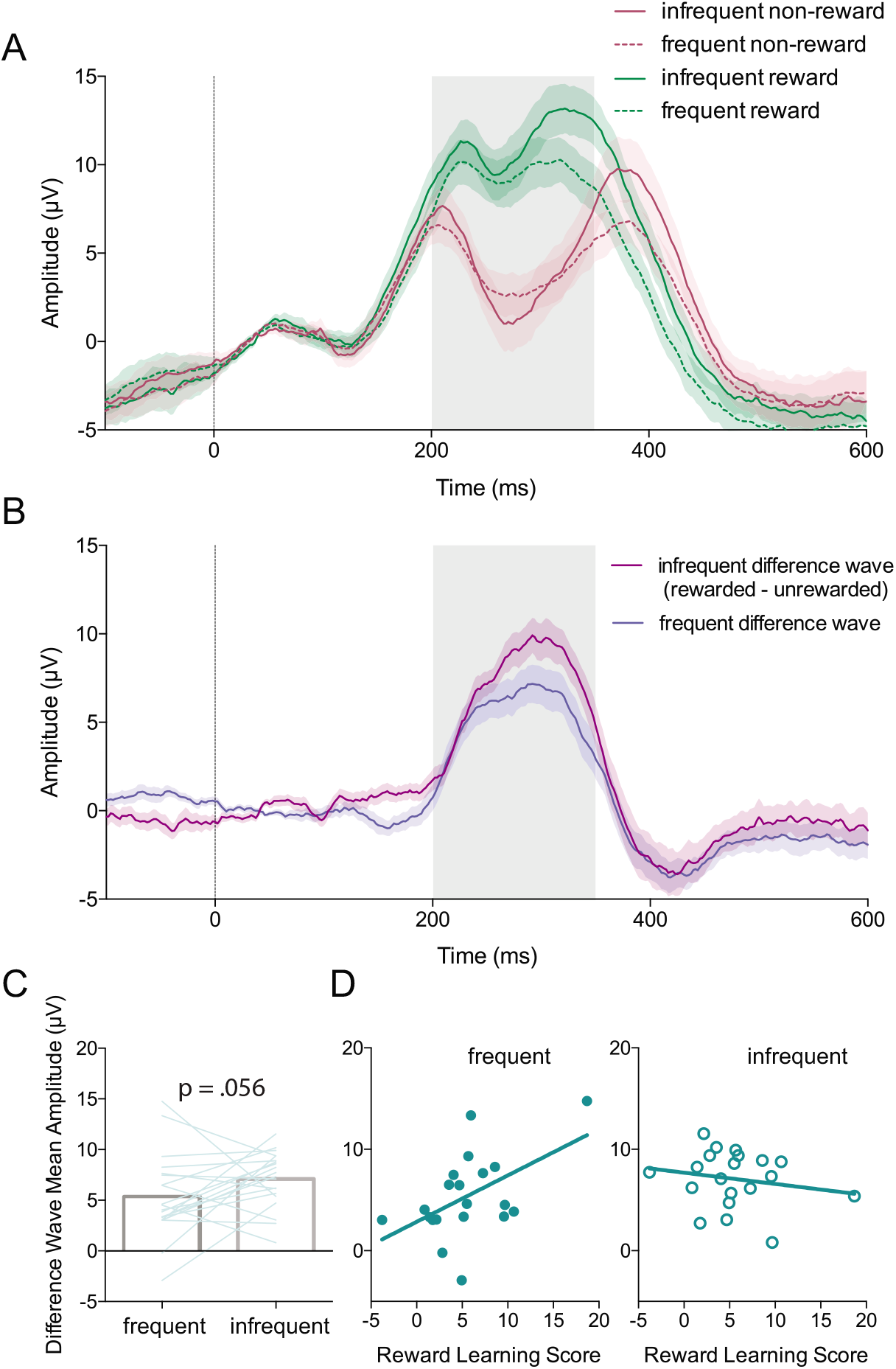
The feedback related negativity was elicited by reward feedback during the reward learning task. A, Trial averaged ERPs recorded from electrode FCz aligned to feedback presentation (0 ms: vertical blue line). Frequent and infrequent reward reflects reward feedback in the high and low reward frequency conditions, respectively. Frequent and infrequent non-reward refers to non-reward feedback in the low and high reward frequency conditions, respectively. Shaded regions: ± SEM. The grey shaded box indicates the temporal window of the FRN. B, The difference waves (reward ERP - non-reward ERP) for frequent and infrequent feedback aligned to feedback presentation. C, The mean amplitude of the difference wave (reward ERP - non-reward ERP) between 200-350ms for infrequent and frequent feedback. D, The mean amplitudes of the difference waves are predictive of behavioral learning scores across participants for the frequent feedback, but not the infrequent feedback.

The mean amplitude of the “infrequent” difference wave was not reliably larger than the mean amplitude of the “frequent” difference wave, although the difference showed a nearly significant trend in this direction (t(19)=1.66, p=.056, paired t-test, one-tailed; Fig 5c). Robust multiple linear regression was used to predict reward learning scores, which we calculated for each subject as the difference in average reach angle between the clockwise and counterclockwise learning conditions, based on the mean values of the “frequent” and “infrequent” difference waves as predictors. The predictors were not correlated (r=0.11, p=.642). The overall multiple regression model was not significant (F(2,17) = 2.72, p = 0.095), with an *R*^2^ of 0.242. Participants’ predicted reward learning score, in degrees of reach angle, is equal to 5.243 + 0.525*β*_1_ − 0.382*β*_2_, where *β*_1_ is the mean value of the “frequent” difference wave in μV, and *β_2_* is the mean value of the “infrequent” difference wave in μV. The “frequent” difference wave was a significant predictor of the reward learning score (t(17) = 2.17, p = 0.044), while the “infrequent” difference wave was not a significant predictor of the reward learning score (t(17) = −1.06, p = 0.30). Figure 5d shows the relationships between the “frequent” and “infrequent” FRN amplitudes and the reward learning score.

#### Visuomotor rotation task

Figure 6a shows the event related potentials (ERPs) recorded from electrode FCz during the visuomotor rotation condition, averaged across participants. The mean value of the “small VMR” difference wave recorded from FCz between 200-350 ms was not significantly different from zero (M=−0.21 *μ*V, SD=1.29, Z=−0.67, W = 87, p=0.50, Wilcoxon signed-ranks test, Fig. 6c). Similarly, the mean value of the “large VMR” difference wave recorded from FCz between 200-350 ms was not significantly different than zero (M=−0.26 *μ*V, SD=1.22, t(19)=−0.97, p=0.34, one-sample t-test, Fig. 6c). These findings indicate that the VMRs imposed in the visuomotor rotation task did not reliably elicit a FRN.

**Figure 6:**
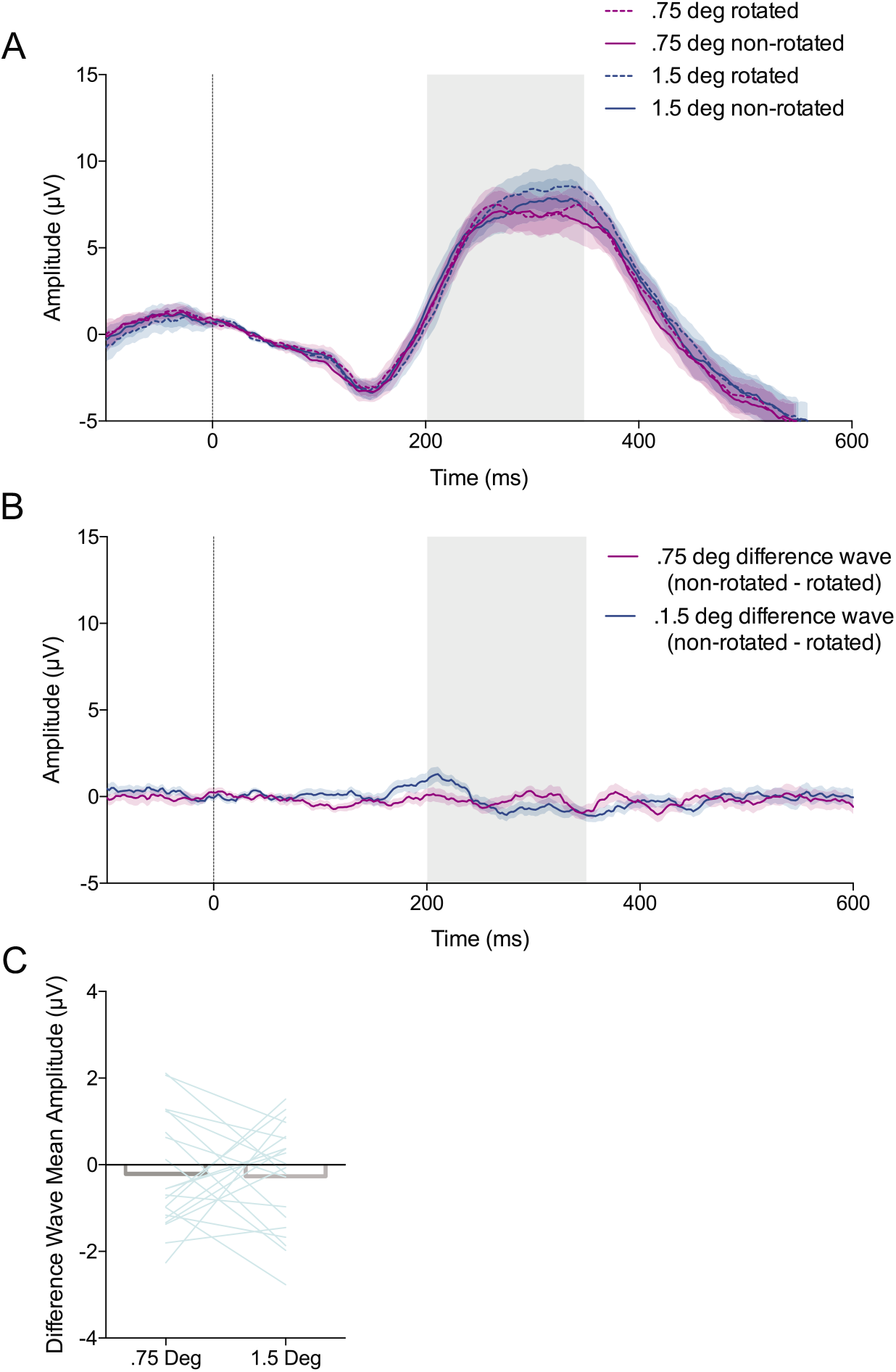
The feedback related negativity was not elicited by sensory error feedback during the visuomotor rotation task. A, Trial averaged ERPs recorded from electrode FCz aligned to feedback presentation (0 ms: vertical line). Shaded regions: ± SEM. The grey shaded box indicates the temporal window of the FRN. B, The difference waves non rotated ERP - rotated ERP) for the .75 and 1.5 degree rotation conditions aligned to feedback presentation. C, The mean amplitude of the difference wave (non rotated ERP - rotated ERP) between 200-350ms for the .75 and 1.5 degree rotation condition (Error bars: ± SEM).

The mean value of the “endpoint error” difference wave recorded from FCz between 200-350 ms was not significantly different than zero (M=0.61 *μ*V, SD=3.28, t(19)=0.82, p=.42, one-sample t-test), indicating that a FRN did not reliably occur on the basis of endpoint error feedback. The fact that we were able to detect a FRN in the reward learning task but not in the visuomotor rotation task is consistent with the notion that the FRN reflects reward processing but not sensory error processing, and that our experimental design successfully dissociated the two.

### P300 Results

#### Reward learning task

Figure 7A shows event related potentials (ERPs) recorded from electrode Pz during the reward learning condition, averaged across participants. We performed a 2×2 repeated measures ANOVA on P300 amplitude with factors expectancy and reward. Figure 7b shows P300 amplitude for each condition, averaged across participants We found a significant main effect of feedback expectancy (F(1,19)=97.16, p<.001), indicating that P300 amplitude was significantly larger in the infrequent feedback conditions. We also found a significant main effect of reward, (F(1,19)=13.18, p=.002), indicating P300 amplitude was larger following rewarded trials compared to unrewarded trials. We found no reliable interaction between reward and expectancy, (F(1,19)=0.992, p=.332). P300 amplitude was not significantly correlated to reward learning scores for any of the four feedback conditions: frequent reward (*R*^2^=0.17, F(1,18)=0.97, p=0.34), infrequent reward (*R*^2^=0.030, F(1,18)=0.558, p=0.47), frequent non-reward (*R*^2^=0.067, F(1,18)=1.3, p=0.27), and infrequent non-reward (*R*^2^=0.06, F(1,18)=1.15, p=0.30).

**Figure 7:**
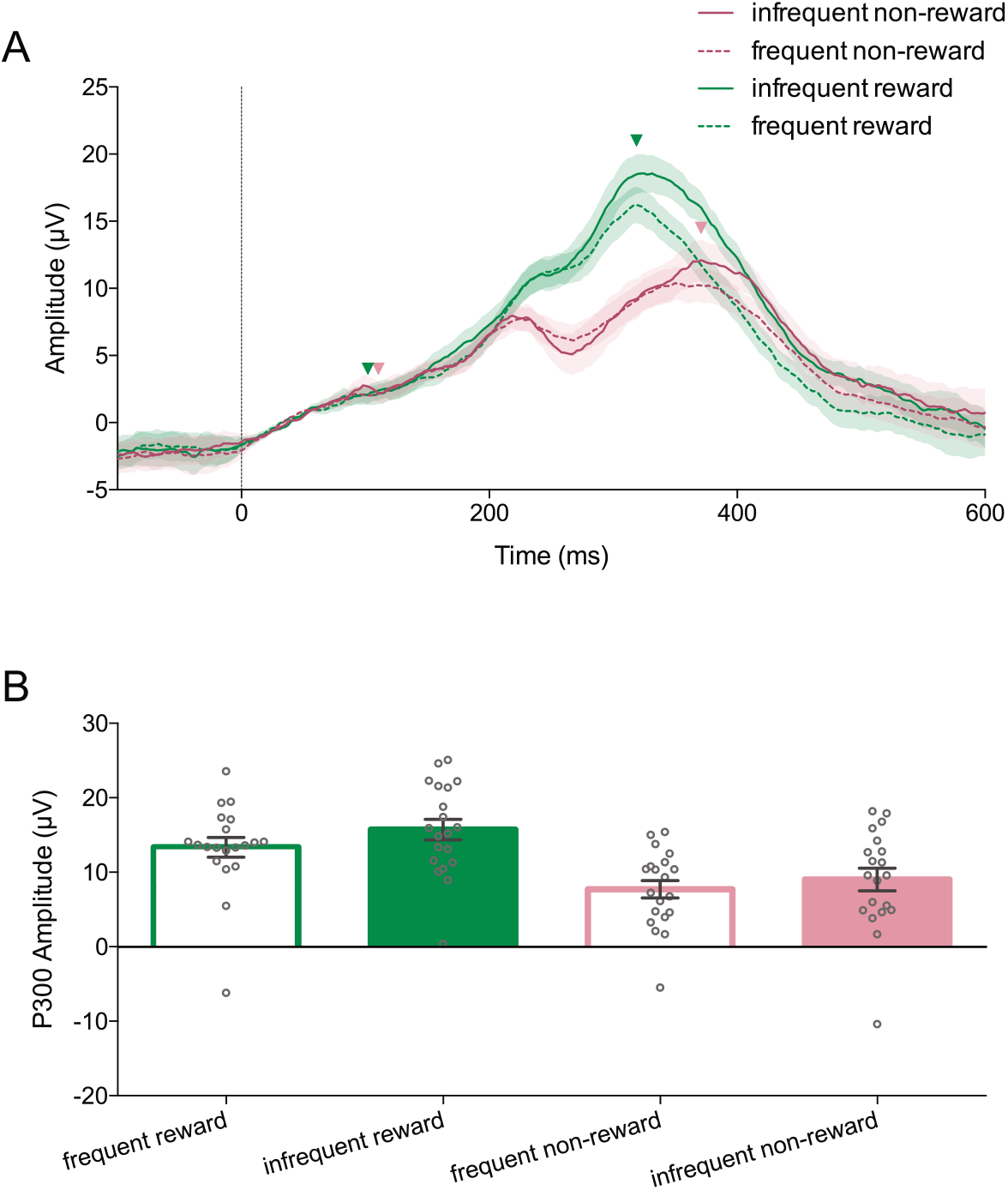
The P300 is modulated by feedback during the reward learning task. A, Trial averaged ERPs recorded from electrode Pz aligned to feedback presentation (0 ms: vertical line). Shaded regions: ± SEM. Arrows indicate the time points for the base and peak of the P300. B, P300 amplitude in each feedback condition (Error bars: ± SEM). P300 amplitude is larger for rewarded feedback relative to unrewarded feedback, and infrequent feedback relative to frequent feedback.

#### Visuomotor rotation task

Figure 8a shows event related potentials (ERPs) recorded from electrode Pz during the visuomotor rotation task, averaged across participants. We first-tested for an effect of the visuomotor rotation on P300 amplitude by comparing non-rotated feedback trials and rotated feedback trials. We performed a two-way repeated measures ANOVA with factors presence of rotation and size of rotation (Figure 8b). We did not find significant main effects of presence of rotation (F(1,19)=2.917, p=.104). We also did not find a main effect of size of rotation (F(1,19)=3.087, p=.095). We did find a significant interaction effect between presence of rotation and rotation magnitude (F(1,19)=8.728, p=.008). We performed planned pairwise comparisons using Bonferroni corrected t-tests between non-rotated and rotated conditions separately for the small and large error conditions. We found that P300 amplitude was significantly greater for rotated, compared to non-rotated, feedback in the 1.5 deg rotation condition (t(19)= 2.83, p=0.021, Bonferroni corrected), but not the .75 deg rotation condition (t(19)=0.09, p=0.93).

**Figure 8:**
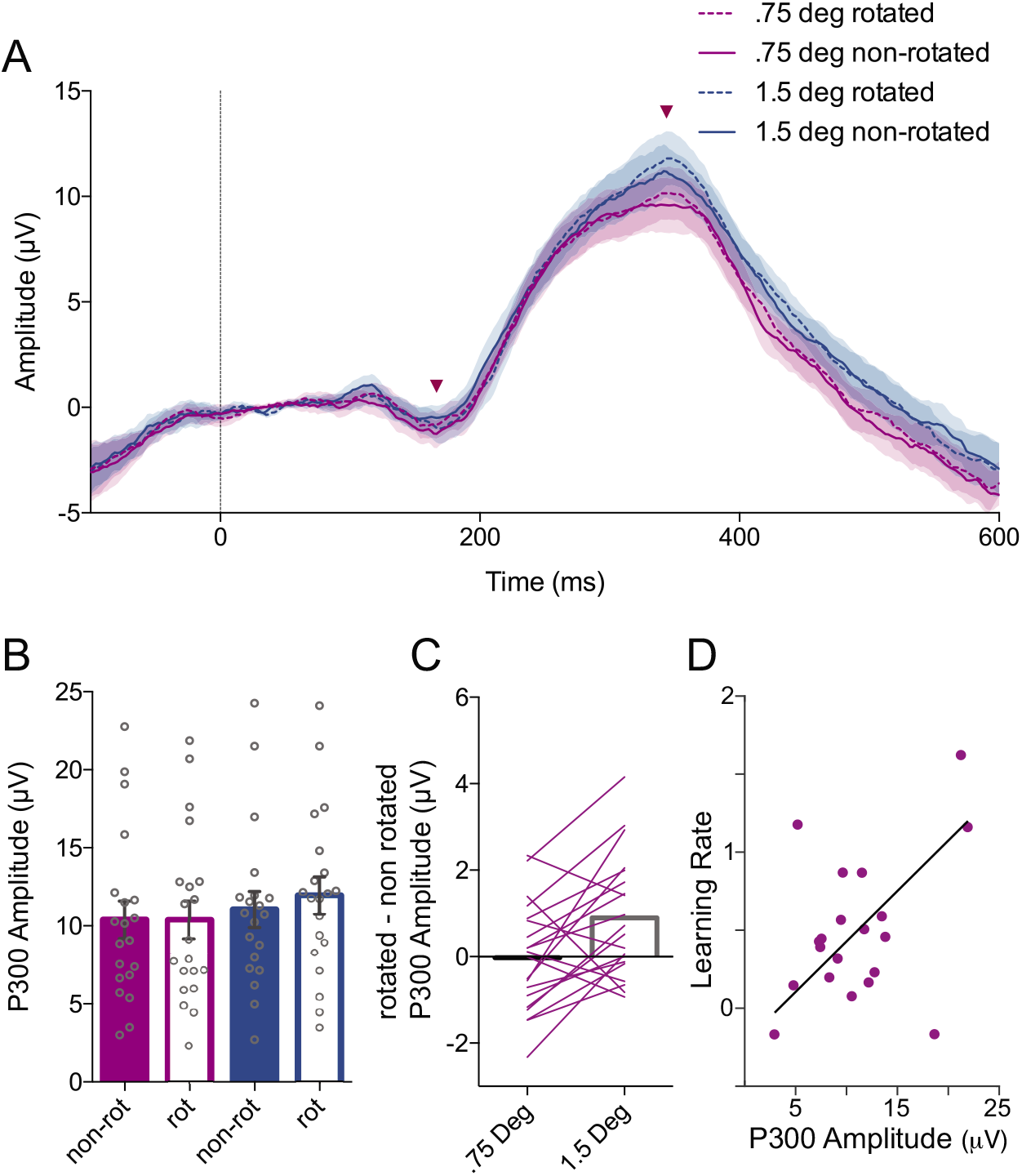
The P300 reflects sensory error processing during the visuomotor rotation task. A, Trial average event related potentials (ERPs) recorded from electrode Pz aligned to feedback presentation (0 ms). Shaded regions: ± SEM. Arrows indicate the time points for the base and peak of the P300. B, The peak-to-peak amplitude of the P300 during the visuomotor rotation task (Error bars: ± SEM). C, P300 amplitude was larger for rotated than non-rotated trials in the 1.5 deg rotation condition but not the .75 deg rotation condition. D, P300 amplitude during adaptation was predicted learning rate. Line of best fit corresponds to robust linear regression using iteratively reweighted least squares.

Robust linear regression was used to predict behavioural learning rate in the visuomotor rotation task based on the amplitude of the P300 measured in the “adaptation” condition ERPs (Fig 8d). P300 amplitude was a significant predictor of learning rate (F(1,18) = 15.9, p < 0.0001), with an *R*^2^ of 0.469. Participants’ predicted learning rate is equal to −0.219 + 0.065*β*_1_, where *β*_1_ is the P300 amplitude in microvolts.

## Discussion

We observed neural correlates of reward and sensory error feedback processing during motor adaptation. We employed reaching tasks that were designed to isolate reward and sensory error based learning while producing comparable changes in reach angle. In both tasks, learning occurred in response to discrete feedback shown after each movement. By examining event related potentials (ERPs) elicited by feedback delivered at the end of movement we avoided potential confounds caused by neural activity or artifacts related to movement execution, motion of the limb, and online error correction. We observed that the feedback related negativity (FRN) was elicited by binary reward feedback but was not by sensory error feedback. This suggests that the process generating the FRN is not necessary for sensory error based learning, and supports the idea that the processes underlying the FRN are specific to reinforcement learning. The P300 occurred in response to both reinforcement and sensory error feedback, and P300 amplitude was modulated by visuomotor rotation, reward, and surprise. In the visuomotor rotation task, P300 amplitude depended on the size of the visuomotor perturbation, and was correlated to learning rate across participants. This suggests that the P300 might reflect general processing of prediction error that is particularly important for sensory error based motor adaptation.

### The FRN Reflects Processing of Reward Feedback but not Sensory Error Feedback

Although motor adaptation has traditionally focused on sensory error learning, recent work suggests that reinforcement learning processes can also contribute to motor adaptation. In the present study, reward based learning was isolated from sensory error based learning during the reward learning task by providing only binary reward feedback in the absence of visual information concerning the position of the hand relative to the target. This feedback elicited a well characterized fronto-central event related potential (ERP) component known as the FRN.

In reinforcement learning theory, agents estimate the expected value of reward outcomes associated with actions, and actions are selected according to predictions regarding the value of reward outcomes. Reward prediction error (RPE) is the difference between the observed value and the predicted value of a reward outcome. Estimates of expected value are updated proportionally to RPE. The true expected value of a reward outcomes is equal to the product of reward magnitude and the probability of reward. In our reward learning task the reward magnitude was fixed, and thus can be represented as a binary quantity (0 or 1), and so the expected value of a particular action on each trial (e.g. the direction of hand movement) was directly proportional to the probability of reward. Therefore, reward feedback should elicit a positive RPE with increasing magnitude when reward is less probable. Conversely, non-reward should elicit negative RPE with increasing magnitude when reward is more probable.

The FRN was observed in the reward learning task as a difference in voltage between ERPs elicited by non-reward feedback and those elicited by reward feedback. A large body of literature has shown that the FRN is larger for infrequent outcomes than frequent outcomes, which supports the theory that the FRN encodes a signed RPE (Cohen et al. 2007; Holroyd and Krigolson 2007; Eppinger et al. 2008; Holroyd et al. 2011; Kreussel et al. 2012; Walsh and Anderson 2012). In the present study, the FRN was larger for improbable feedback than for probable feedback, although the difference was not statistically reliable (p=.056). This result is potentially due to the relatively small difference in reward frequency experienced between the low and high reward frequency conditions (38.6% and 67.0%, respectively) when compared to other studies. We decided to avoid using very low or very high reward frequency as we found it to produce highly variable and strategic behavior in the task.

A prominent theory proposes that the FRN reflects activity originating in the anterior cingulate cortex driven by phasic dopaminergic signaling of RPE (Holroyd and Coles 2002; Santesso et al. 2009). This activity is purported to underlie reward-based learning processes by integrating RPE to affect future action selection. We sought to examine the role of the FRN in learning by testing the correlation between the amplitude of the FRN and the extent of learning as assessed behaviorally.

In typical reward learning paradigms, successful learning is correlated with increased reward frequency, and as such the relationship between the FRN and the magnitude of learning is confounded by any effect of reward frequency on FRN amplitude. In the current study, the reward learning task was adaptive to learning such that the overall reward frequency was largely decoupled from the extent of adaptation. We found that the extent of behavioral learning was predicted by the amplitude of the FRN for the frequent feedback condition but not the infrequent feedback condition. One possible explanation for this discrepancy is that participants responded reliably to large RPEs elicited by infrequent feedback as they were most salient, but the limiting factor in learning was a sensitivity to small RPEs. Even though small RPEs should produce less learning per trial than large RPEs, in our task they were by definition more numerous and as such could contribute comparably to the overall extent of learning. Another possibility is that the measure of FRN amplitude became less reliable in the infrequent feedback condition due to interference by surprise related signals that were not directly related to RL, such as the P300. Relationships between the FRN and learning have been observed for tasks such as time estimation and discrete motor sequence learning (Holroyd and Krigolson 2007; Helden et al. 2010). As such, the present findings support the idea that the same RPE processing mechanism can drive learning across both cognitive and motor tasks. However, correlations with limited sample sizes can be unreliable, and the relationship between FRN amplitude and reward learning in the present study was not particularly strong. Future work is needed to further test this relationship, preferably through causal manipulation and behavioural tasks that produce less variable and idiosyncratic learning behavior.

The sensory error-based learning task allowed us to dissociate the neural signatures of reward based learning and sensory error based learning. Sensory error often coincides with task error and reward omission, as failure to achieve the expected sensory consequences of a motor command usually entails failure to achieve the subjective goal of the task. Achieving task goals may be inherently rewarding or explicitly rewarded, as in the current study. Our visuomotor rotation task was designed to elicit learning through sensory error feedback while minimizing task error, as the perturbations were relatively small and rarely resulted in cursor feedback outside of the target. We did not observe the presence of the FRN in response to sensory error feedback, despite reliable observation of behavioural adaptation and the P300 neural response. This suggests that the process which generates the FRN is not necessary for adaptation to sensory errors.

Our results suggest that the ERP responses to reward and sensory error processing can be dissociated in motor adaptation, and that the FRN is specific to reward based learning processes. Although the FRN is classically associated with reinforcement learning, recent work has identified the FRN or the closely related error-related negativity (ERN) in various motor learning and execution tasks involving sensory error signals. These studies either concluded that reinforcement and sensory error based learning processes share common neural resources, or they simply do not distinguish between these two processes (Krigolson et al. 2008; Torrecillos et al. 2014; MacLean et al. 2015). We argue that the brain processes reward and sensory error feedback through distinct mechanisms, but that the two processes can be confounded when perturbations causing sensory error are also evaluated as an implicit failure to meet task goals. In this case, the brain could process a reward and sensory prediction error independently, although learning might be driven primarily by the sensory prediction error. This is consistent with behavioral studies showing that sensory error based learning is the primary driver of behavioral change when both reward and sensory feedback is provided (Izawa and Shadmehr 2011; Cashaback et al. 2017).

Recently, Savoie et al. (2018) also examined the effects of sensory error on EEG responses while carefully controlling for reward or task related errors. Following Mazzoni and Krakauer (2006), a 45 deg visuomotor rotation was imposed on continuous cursor feedback, and participants were instructed to reach to a second target opposite to the rotation. In this paradigm, participants do not experience failure to achieve task goals or reward, as they effectively counteract the rotation through strategic aiming. Nonetheless, participants automatically and implicitly adapt to the sensory error caused by the rotation. This strategic condition was contrasted to a condition with no instructed strategy, in which participants had already adapted fully to a 45 deg rotation, and thus experienced no task or sensory error. Unlike the current study, the authors report a prominent mid-frontal negativity resembling the FRN in the strategic condition, despite a lack of task or reward related errors. One possible explanation for these conflicting results is that the FRN can be elicited by sensory error, but that the visuomotor rotations used in the current study were simply too small to elicit an observable FRN. Another possible explanation is that the response observed by Savoie et al. was not an FRN elicited by failure to achieve reward or task goals, but another frontal negativity related to implementation of the strategy such as the N200, which can be indistinguishable from the FRN (Holroyd et al. 2008). The N200 is elicited by response conflict and cognitive control, which may have occurred during the strategic aiming condition as participants were required to inhibit the prepotent response of reaching directly towards the target and instead implement the strategic aiming response (Nieuwenhuis et al. 2003; Folstein and Van Petten 2008; Enriquez-Geppert et al. 2010).

### The P300 is Modulated by Sensory Error, Surprise, and Reward

During the visuomotor rotation task in the present study, we observed a P300 event related potential (ERP) in response to reach endpoint position feedback, and we found that P300 amplitude was sensitive to the magnitude of sensory error. P300 amplitude was increased by the larger but not the smaller visuomotor rotation. Learning in visuomotor rotation paradigms is thought to be driven primarily by sensory error based learning, and as such our findings suggest that the P300 observed in this task might reflect neural activity that is related to processing of sensory error underlying motor adaptation (Izawa and Shadmehr 2011).

It is important to note that a P300 response is typically elicited by stimulus processing in tasks unrelated to sensory error-based motor adaptation, including the reward learning task in the present study. A prominent and enduring theory proposes that the P300 reflects cortical processing related to the updating of a neural model of stimulus context upon processing of unexpected stimuli (Donchin 1981; Donchin and Coles 1988; Polich 2007). This theory resembles an account of sensory error based motor adaptation in which the updating of an internal model of motor dynamics occurs when sensory input differs from the predictions of the internal model (Wolpert et al. 1995; Synofzik et al. 2008). It is possible that the P300 reflects a general aspect of prediction error processing that is common to both sensorimotor and cognitive function. The P300 is typically localized to parietal regions, which are implicated in processing sensory prediction error during adaptation to visuomotor rotation (Bledowski et al. 2004; Diedrichsen et al. 2005; Linden 2005; Tanaka et al. 2009). Consistent with the P300 underlying sensory prediction error processing, cerebellar damage impairs sensory error based adaptation and results in P300 abnormalities (Tachibana et al. 1995; Martin et al. 1996; Maschke et al. 2004; Paulus et al. 2004; Smith and Shadmehr 2005; Mannarelli et al. 2015; Therrien et al. 2015).

Previous studies have examined P300 responses elicited by sensory error feedback. It has been demonstrated that P300 amplitude decreases along with the magnitude of reach errors during the course of prism adaptation (MacLean et al. 2015). This suggests that the P300 is modulated by sensory error, but it does not rule out the possibility that the P300 simply attenuated with time. The P300 has also been reported to occur in response to random shifts in target location (Krigolson et al. 2008), another type of visuo-spatial error which, however, does not reliably produce learning (Diedrichsen et al. 2005). Both of these studies also reported FRN components occurring in response to task errors. In the present study a P300 response was observed that is correlated to behavioral learning and isolated from the FRN.

In our reward learning task, P300 amplitude was modulated by feedback valence and expectancy but was not correlated to learning. The finding that P300 shows a larger positive amplitude for infrequent feedback regardless of valence is consistent with previous reports and supports the notion that the P300 reflects a general prediction error processing when the stimulus differs from expectation (Hajcak et al. 2005, 2007; Wu and Zhou 2009; Pfabigan et al. 2011). This is distinct from the interaction between valence and expectancy that is characteristic of encoding reward prediction error, in which surprise has opposite effects on the response to reward and non reward. We also found that the P300 was larger for reward than non reward feedback. Some previous studies have shown a similar effect of reward valence on P300 amplitude (Hajcak et al. 2005; Wu and Zhou 2009; Leng and Zhou 2010; Zhou et al. 2010), while others have not (Yeung and Sanfey 2004; Sato et al. 2005; Pfabigan et al. 2011). This finding is also consistent with idea that the P300 reflects updating of a model of stimulus context in response to prediction error, as reward feedback would indicate that the previous action was rewarding while all other possible actions were not rewarding. Non reward feedback would only indicate that the previous action was not rewarding, and thus carries less information for updating of the internal representation of the task.

### Outstanding Questions

In visuomotor rotation paradigms, a dissociation has been drawn between explicit and implicit learning processes which contribute to learning in a largely independent manner (Mazzoni and Krakauer 2006; Krakauer 2009; Benson et al. 2011; Heuer and Hegele 2011; Taylor et al. 2014; McDougle et al. 2015, 2016; Schween and Hegele 2017). The implicit process occurs automatically, without conscious awareness, and constitutes a recalibration of the visuomotor mapping. The explicit process is characterized by conscious and strategic changes in aiming intended to counteract experimental perturbations. The implicit component is known to be dependent on cerebellar processes, while explicit learning may rely on prefrontal and premotor cortex (Taylor et al. 2010; Heuer and Hegele 2011; Taylor and Ivry 2014; McDougle et al. 2016).

In the current study, relatively small perturbations of feedback produce gradual changes in reach direction. This gradual form of adaptation is thought to primarily recruit the implicit adaptation process (Klassen et al. 2005; Michel et al. 2007; Saijo and Gomi 2010). Nonetheless, it is possible that a mixture of implicit and strategic learning contributes the observed adaptation, especially considering the finding that visual feedback restricted to movement endpoint elicits less implicit learning relative to continuous feedback, and that strategic aiming is employed to reduce residual error (Taylor et al. 2014). Further work is necessary to determine whether the neural generators of the P300 observed in the VMR task contribute specifically to implicit or strategic learning processes.

Similarly, it is not clear whether adaptation in the RL task occurred implicitly or through strategic processes. The extent of learning was variable and idiosyncratic, which may reflect differences in awareness of the manipulation or conscious strategy (Holland et al. 2018). Recent work has shown that when participants learn to produce reach angles directed away from a visual target through binary reward feedback, adaptation is dramatically reduced by instructions to cease any strategic aiming, suggesting a dominant explicit component to reward based reach adaptation (Codol et al. 2018; Holland et al. 2018). Nonetheless, after learning a 25 degree rotation through binary feedback and being instructed to cease strategic aiming, small changes in reach angle persist (Holland et al. 2018). It is not clear whether this residual adaptation can be attributed to an implicit form of reward learning or whether it reflects use dependent plasticity, but it suggests that implicit reward learning may occur for small changes in reach angle, such as those observed in the present study. Future work should determine whether the FRN and P300 are specifically related to strategic or explicit reward based motor adaptation, especially considering evidence from sequence and cognitive learning domains that the FRN relates more closely to explicit processes (Rüsseler et al. 2003; Loonis et al. 2017).

## References

Alexander WH, Brown JW. Medial prefrontal cortex as an action-outcome predictor. Nat Neurosci 14: 1338–1344, 2011.

Baker TE, Holroyd CB. Dissociated roles of the anterior cingulate cortex in reward and conflict processing as revealed by the feedback error-related negativity and N200. Biol Psychol 87: 25–34, 2011a.

Baker TE, Holroyd CB. Dissociated roles of the anterior cingulate cortex in reward and conflict processing as revealed by the feedback error-related negativity and N200. Biol Psychol 87: 25–34, 2011b.

Becker MPI, Nitsch AM, Miltner WHR, Straube T. A single-trial estimation of the feedback-related negativity and its relation to BOLD responses in a time-estimation task. J Neurosci 34: 3005–3012, 2014.

Benson BL, Anguera JA, Seidler RD. A spatial explicit strategy reduces error but interferes with sensorimotor adaptation. Journal of neurophysiology 105: 2843–2851, 2011.

Bédard P, Sanes JN. Brain representations for acquiring and recalling visual-motor adaptations. Neuroimage 101: 225–235, 2014.

Bledowski C, Prvulovic D, Hoechstetter K, Scherg M, Wibral M, Goebel R, Linden DEJ. Localizing P300 generators in visual target and distractor processing: A combined event-related potential and functional magnetic resonance imaging study. J Neurosci 24: 9353–9360, 2004.

Botvinick Y. Reinforcement learning, conflict monitoring, and cognitive control: An integrative model of cingulate-striatal interactions and the ERN. Neural basis of motivational and cognitive control.

Brooks JL, Zoumpoulaki A, Bowman H. Data-driven region-of-interest selection without inflating Type I error rate. Psychophysiology 54: 100–113, 2017.

Carlson JM, Foti D, Mujica-Parodi LR, Harmon-Jones E, Hajcak G. Ventral striatal and medial prefrontal BOLD activation is correlated with reward-related electrocortical activity: A combined ERP and fMRI study. Neuroimage 57: 1608–1616, 2011.

Cashaback JGA, McGregor HR, Mohatarem A, Gribble PL. Dissociating error-based and reinforcement-based loss functions during sensorimotor learning. PLoS Comput Biol 13: e1005623, 2017.

Codol O, Holland PJ, Galea JM. The relationship between reinforcement and explicit control during visuomotor adaptation. Sci Rep 8: 9121, 2018.

Cohen MX, Elger CE, Ranganath C. Reward expectation modulates feedback-related negativity and EEG spectra. Neuroimage 35: 968–978, 2007.

Delorme A, Makeig S. EEGLAB: An open source toolbox for analysis of single-trial EEG dynamics including independent component analysis. J Neurosci Methods 134: 9–21, 2004.

Diedrichsen J, Hashambhoy Y, Rane T, Shadmehr R. Neural correlates of reach errors. J Neurosci 25: 9919–9931, 2005.

Donchin E. Surprise!… surprise? Psychophysiology 18: 493–513, 1981.

Donchin E, Coles MGH. Is the P300 component a manifestation of context updating? Behav Brain Sci 11: 357–374, 1988.

Enriquez-Geppert S, Konrad C, Pantev C, Huster RJ. Conflict and inhibition differentially affect the n200/p300 complex in a combined go/nogo and stop-signal task. Neuroimage 51: 877–887, 2010.

Eppinger B, Kray J, Mock B, Mecklinger A. Better or worse than expected? Aging, learning, and the ERN. Neuropsychologia 46: 521–539, 2008.

Fabiani M, Gratton G, Karis D, Donchin E, Others. Definition, identification, and reliability of measurement of the P300 component of the event-related brain potential. Advances in psychophysiology 2: 78, 1987.

Folstein JR, Van Petten C. Influence of cognitive control and mismatch on the n2 component of the erp: A review. Psychophysiology 45: 152–170, 2008.

Galea JM, Mallia E, Rothwell J, Diedrichsen J. The dissociable effects of punishment and reward on motor learning. Nat Neurosci 18: 597–602, 2015.

Galea JM, Ruge D, Buijink A, Bestmann S, Rothwell JC. Punishment-Induced behavioral and neurophysiological variability reveals Dopamine-Dependent selection of kinematic movement parameters. J Neurosci 33: 3981–3988, 2013.

Glimcher PW. Understanding dopamine and reinforcement learning: The dopamine reward prediction error hypothesis. Proc Natl Acad Sci U S A 108: 15647–15654, 2011.

Hajcak G, Holroyd CB, Moser JS, Simons RF. Brain potentials associated with expected and unexpected good and bad outcomes. Psychophysiology 42: 161–170, 2005.

Hajcak G, Moser JS, Holroyd CB, Simons RF. It’s worse than you thought: The feedback negativity and violations of reward prediction in gambling tasks. Psychophysiology 44: 905–912, 2007.

HeldenJ van der, Boksem MAS, Blom JHG. The importance of failure: Feedback-related negativity predicts motor learning efficiency. Cereb Cortex 20: 1596–1603, 2010.

Heuer H, Hegele M. Generalization of implicit and explicit adjustments to visuomotor rotations across the workspace in younger and older adults. J Neurophysiol 106: 2078–2085, 2011.

Heydari S, Holroyd CB. Reward positivity: Reward prediction error or salience prediction error? Psychophysiology 53: 1185–1192, 2016.

Holland P, Codol O, Galea JM. Contribution of explicit processes to reinforcement-based motor learning. J Neurophysiol 119: 2241–2255, 2018.

Holroyd CB, Coles MGH. The neural basis of human error processing: Reinforcement learning, dopamine, and the error-related negativity. Psychol Rev 109: 679–709, 2002.

Holroyd CB, Krigolson OE. Reward prediction error signals associated with a modified time estimation task. Psychophysiology 44: 913–917, 2007.

Holroyd CB, Krigolson OE, Lee S. Reward positivity elicited by predictive cues. Neuroreport 22: 249–252, 2011.

Holroyd CB, Pakzad-Vaezi KL, Krigolson OE. The feedback correct-related positivity: Sensitivity of the event-related brain potential to unexpected positive feedback. Psychophysiology 45: 688–697, 2008.

Huang VS, Haith A, Mazzoni P, Krakauer JW. Rethinking motor learning and savings in adaptation paradigms: Model-free memory for successful actions combines with internal models. Neuron 70: 787–801, 2011.

Inoue K, Kawashima R, Satoh K, Kinomura S, Sugiura M, Goto R, Ito M, Fukuda H. A PET study of visuomotor learning under optical rotation. Neuroimage 11: 505–516, 2000.

Inoue M, Uchimura M, Kitazawa S. Error signals in motor cortices drive adaptation in reaching. Neuron 90: 1114–1126, 2016.

Izawa J, Shadmehr R. Learning from sensory and reward prediction errors during motor adaptation. PLoS Comput Biol 7: e1002012, 2011.

Klassen J, Tong C, Flanagan JR. Learning and recall of incremental kinematic and dynamic sensorimotor transformations. Exp Brain Res 164: 250–259, 2005.

Krakauer JW. Motor learning and consolidation: The case of visuomotor rotation. In: Progress in motor control. Springer, 2009, p. 405–421.

Krakauer JW, Ghilardi M-F, Mentis M, Barnes A, Veytsman M, Eidelberg D, Ghez C. Differential cortical and subcortical activations in learning rotations and gains for reaching: A PET study. J Neurophysiol 91: 924–933, 2004.

Kreussel L, Hewig J, Kretschmer N, Hecht H, Coles MGH, Miltner WHR. The influence of the magnitude, probability, and valence of potential wins and losses on the amplitude of the feedback negativity. Psychophysiology 49: 207–219, 2012.

Krigolson OE, Holroyd CB, Van Gyn G, Heath M. Electroencephalographic correlates of target and outcome errors. Exp Brain Res 190: 401–411, 2008.

Leng Y, Zhou X. Modulation of the brain activity in outcome evaluation by interpersonal relationship: An ERP study. Neuropsychologia 48: 448–455, 2010.

Linden DEJ. The P300: Where in the brain is it produced and what does it tell us? Neuroscientist 11: 563–576, 2005.

Loonis RF, Brincat SL, Antzoulatos EG, Miller EK. A Meta-Analysis suggests different neural correlates for implicit and explicit learning. Neuron 96: 521–534.e7, 2017.

MacLean SJ, Hassall CD, Ishigami Y, Krigolson OE, Eskes GA. Using brain potentials to understand prism adaptation: The error-related negativity and the P300. Front Hum Neurosci 9, 2015.

Mannarelli D, Pauletti C, De Lucia MC, Currà A, Fattapposta F. Insights from ERPs into attention during recovery after cerebellar stroke: A case report. Neurocase 21: 721–726, 2015.

Marko MK, Haith AM, Harran MD, Shadmehr R. Sensitivity to prediction error in reach adaptation. J Neurophysiol 108: 1752–1763, 2012.

Martin TA, Keating JG, Goodkin HP, Bastian AJ, Thach WT. Throwing while looking through prisms. I. Focal olivocerebellar lesions impair adaptation. Brain 119 (Pt 4): 1183–1198, 1996.

Maschke M, Gomez CM, Ebner TJ, Konczak J. Hereditary cerebellar ataxia progressively impairs force adaptation during goal-directed arm movements. J Neurophysiol 91: 230–238, 2004.

Mazzoni P, Krakauer JW. An implicit plan overrides an explicit strategy during visuomotor adaptation. J Neurosci 26: 3642–3645, 2006.

McDougle SD, Bond KM, Taylor JA. Explicit and implicit processes constitute the fast and slow processes of sensorimotor learning. Journal of Neuroscience 35: 9568–9579, 2015.

McDougle SD, Ivry RB, Taylor JA. Taking aim at the cognitive side of learning in sensorimotor adaptation tasks. Trends Cogn Sci 20: 535–544, 2016.

Michel C, Pisella L, Prablanc C, Rode G, Rossetti Y. Enhancing visuomotor adaptation by reducing error signals: Single-step (aware) versus multiple-step (unaware) exposure to wedge prisms. J Cogn Neurosci 19: 341–350, 2007.

Miltner WH, Braun CH, Coles MG. Event-related brain potentials following incorrect feedback in a time-estimation task: Evidence for a “generic” neural system for error detection. Journal of cognitive neuroscience 9: 788–798, 1997.

Nieuwenhuis S, Holroyd CB, Mol N, Coles MGH. Reinforcement-related brain potentials from medial frontal cortex: Origins and functional significance. Neurosci Biobehav Rev 28: 441–448, 2004.

Nieuwenhuis S, Yeung N, Van Den Wildenberg W, Ridderinkhof KR. Electrophysiological correlates of anterior cingulate function in a go/no-go task: Effects of response conflict and trial type frequency. Cognitive, affective, & behavioral neuroscience 3: 17–26, 2003.

Nikooyan AA, Ahmed AA. Reward feedback accelerates motor learning. J Neurophysiol 113: 633–646, 2015.

Paulus KS, Magnano I, Conti M, Galistu P, D’Onofrio M, Satta W, Aiello I. Pure post–stroke cerebellar cognitive affective syndrome: A case report. Neurol Sci 25: 220–224, 2004.

Pekny SE, Izawa J, Shadmehr R. Reward-dependent modulation of movement variability. Journal of Neuroscience 35: 4015–4024, 2015.

Pfabigan DM, Alexopoulos J, Bauer H, Sailer U. Manipulation of feedback expectancy and valence induces negative and positive reward prediction error signals manifest in event-related brain potentials. Psychophysiology 48: 656–664, 2011.

Polich J. Updating P300: An integrative theory of P3a and P3b. Clin Neurophysiol 118: 2128–2148, 2007.

Reuter E-M, Pearcey GE, Carroll TJ. Greater neural responses to trajectory errors are associated with superior force field adaptation in older adults. Experimental gerontology 110: 105–117, 2018.

Rüsseler J, Kuhlicke D, Münte TF. Human error monitoring during implicit and explicit learning of a sensorimotor sequence. Neurosci Res 47: 233–240, 2003.

Saijo N, Gomi H. Multiple motor learning strategies in visuomotor rotation. PLoS One 5: e9399, 2010.

Santesso DL, Evins AE, Frank MJ, Schetter EC, Bogdan R, Pizzagalli DA. Single dose of a dopamine agonist impairs reinforcement learning in humans: Evidence from event-related potentials and computational modeling of striatal-cortical function. Hum Brain Mapp 30: 1963–1976, 2009.

Sato A, Yasuda A, Ohira H, Miyawaki K, Nishikawa M, Kumano H, Kuboki T. Effects of value and reward magnitude on feedback negativity and P300. Neuroreport 16: 407–411, 2005.

Savoie F-A, Thénault F, Whittingstall K, Bernier P-M. Visuomotor prediction errors modulate EEG activity over parietal cortex. Sci Rep 8: 12513, 2018.

Schween R, Hegele M. Feedback delay attenuates implicit but facilitates explicit adjustments to a visuomotor rotation. Neurobiology of learning and memory 140: 124–133, 2017.

Shmuelof L, Huang VS, Haith AM, Delnicki RJ, Mazzoni P, Krakauer JW. Overcoming motor “forgetting” through reinforcement of learned actions. J Neurosci 32: 14617–14621a, 2012.

Smith MA, Shadmehr R. Intact ability to learn internal models of arm dynamics in huntington’s disease but not cerebellar degeneration. J Neurophysiol 93: 2809–2821, 2005.

Synofzik M, Lindner A, Thier P. The cerebellum updates predictions about the visual consequences of one’s behavior. Curr Biol 18: 814–818, 2008.

Tachibana H, Aragane K, Sugita M. Event-related potentials in patients with cerebellar degeneration: Electrophysiological evidence for cognitive impairment. Brain Res Cogn Brain Res 2: 173–180, 1995.

Tan H, Jenkinson N, Brown P. Dynamic neural correlates of motor error monitoring and adaptation during trial-to-trial learning. Journal of Neuroscience 34: 5678–5688, 2014.

Tanaka H, Sejnowski TJ, Krakauer JW. Adaptation to visuomotor rotation through interaction between posterior parietal and motor cortical areas. J Neurophysiol 102: 2921–2932, 2009.

Taylor JA, Ivry RB. Chapter 9 - cerebellar and prefrontal cortex contributions to adaptation, strategies, and reinforcement learning. In: Progress in brain research, edited by Ramnani N. Elsevier, 2014, p. 217–253.

Taylor JA, Klemfuss NM, Ivry RB. An explicit strategy prevails when the cerebellum fails to compute movement errors. Cerebellum 9: 580–586, 2010.

Taylor JA, Krakauer JW, Ivry RB. Explicit and implicit contributions to learning in a sensorimotor adaptation task. J Neurosci 34: 3023–3032, 2014.

Therrien AS, Wolpert DM, Bastian AJ. Effective reinforcement learning following cerebellar damage requires a balance between exploration and motor noise. Brain 139: 101–114, 2015.

Thoroughman KA, Shadmehr R. Learning of action through adaptive combination of motor primitives. Nature 407: 742–747, 2000.

Torrecillos F, Albouy P, Brochier T, Malfait N. Does the processing of sensory and reward-prediction errors involve common neural resources? Evidence from a frontocentral negative potential modulated by movement execution errors. J Neurosci 34: 4845–4856, 2014.

Walsh MM, Anderson JR. Learning from experience: Event-related potential correlates of reward processing, neural adaptation, and behavioral choice. Neurosci Biobehav Rev 36: 1870–1884, 2012.

Wolpert DM, Ghahramani Z, Jordan MI. An internal model for sensorimotor integration. Science 269: 1880–1882, 1995.

Wu Y, Zhou X. The P300 and reward valence, magnitude, and expectancy in outcome evaluation. Brain Res 1286: 114–122, 2009.

Yeung N, Sanfey AG. Independent coding of reward magnitude and valence in the human brain. J Neurosci 24: 6258–6264, 2004.

Zhou Z, Yu R, Zhou X. To do or not to do? Action enlarges the FRN and P300 effects in outcome evaluation. Neuropsychologia 48: 3606–3613, 2010.

